# Sample coverage standardization ranks species richness of saproxylic beetles across European tree genera

**DOI:** 10.64898/2026.09.29.755269

**Authors:** Ronja Hausmann, Anne Chao, Romain Angeleri, Christophe Bouget, Antoine Brin, Cristiana Cocciufa, Anika Gossmann, Martin M. Gossner, Elena Haeler, Jonas Hagge, Marit L. Hertlein, Joakim Hjältén, Peter Kriegel, Thibault Lachat, Laurent Larrieu, Anna L. M. Macagno, Oliver Mitesser, Claudio Sbaraglia, Sebastian Seibold, Sebastian Vogel, Sebastian Zarges, Jörg Müller, Simon Thorn

**Author notes:** **Corresponding author:** Ronja Hausmann.

## Abstract

**Aims:** In forest ecosystems, tree species composition and diversity are fundamental drivers of biodiversity and many species are closely associated to specific tree species as hosts. Saproxylic diversity across tree species may be shaped by interactions between ecological and environmental factors, such as resource availability. However, a throughout and unbiased quantification of diversity across tree species is missing. We investigated patterns an drivers of saproxylic beetle diversity across European tree gnera while accounting for differences in sampling effort.

**Location:** Europe

**Methods:** We used a large dataset, of 25 studies, comprising 500,000 deadwood-dependent beetles, representing 1041 saproxylic beetle species, sampled from 29 tree genera. We accounted for sampling effort by estimating diversity using coverage-based standardization across Hill numbers. We than used linear models to identify drivers of saproxylic beetle diversity across tree genera and assessed differences in community composition using Nonmetric Multidimensional Scaling (NMDS). Additionally, the most abundant beetle families were identified for a subset of 14 genera.

**Results:** *Quercus* supported the highest richness of infrequent, frequent, and highly frequent species, while *Fagus* exhibited the highest observed (unstandardized) species richness. Beetle communities were less differentiated when focusing on infrequent species, whereas tree genera were separated when focusing on frequent and highly frequent species. In addition, we found that resource availability and maximum tree height best predicted infrequent beetle diversity, whereas resource availability was the main driver of frequent species. In contrast, the same analyses for unstandardized data indicated postglacial occurrence as an important factor for infrequent species.

**Main conclusions:** Although resource availability supported the *species–area hypothesis*, standardized data revealed higher richness on smaller trees, partly contradicting this pattern. Our results highlight the importance of accounting for sampling bias and promoting tree diversity, particularly oaks, to sustain saproxylic beetle diversity.

## Introduction

Identifying the factors that influence species diversity and its spatial patterns is a central objective of ecological research and fundamental for conservation management. While environmental constraints, such as temperature, rainfall, evapotranspiration, and primary productivity, account for a considerable proportion of the observed variation in diversity (Hawkins et al., 2003), biodiversity observed at the local scale is additionally determined by the interaction of ecological and environmental constraints, such as the availability of resources (Kostylev et al., 2005; Messmer et al., 2011).

Associations between insects and their host plants are widespread in nature, including phytophagous insects, parasites, and parasitoids (Brändle et al., 2008; Ehrlich & Raven, 1964; Jermy, 1984; Kamiya et al., 2014; Thorn et al., 2015). A meta-analysis revealed that population density and geographical range of hosts can predict species richness of associated parasites (Kamiya et al., 2014). In addition, host size, morphological complexity and the postglacial occurrence of a plant species correlate positively with herbivore species richness (Brändle et al., 2008; Strong & Levin, 1979). Several hypotheses have been proposed to explain these patterns: Kennedy & Southwood (1984) originally formulated three hypotheses, which were later expanded by Brändle & Brandl (2001). The *species-area hypothesis* proposes that species richness increases with resource availability (host biomass) and spatial distribution, with individual trees functioning as habitat islands at a local scale. Similarly, also larger trees are expected to host more phytophagous species due to their higher number of microclimatic niches. The *geological-time hypothesis* suggests that the longer a tree species occurs in an area (e.g., postglacial re-colonization in Europe), the more opportunities for possible colonization and specialization by associated species arise. Finally, the *taxonomic-isolation hypothesis* predicts that host shifts occur more frequently between closely related hosts, because phylogenetically related hosts tend to offer similar habitat conditions (Brändle & Brandl, 2001; Kennedy & Southwood, 1984). These general patterns and hypotheses may also apply to other insect groups with strong host associations, such as saproxylic, i.e., deadwood-associated, insects in forest ecosystems.

Saproxylic insects depend on dead or dying wood during part of their life cycle (Speight, 1989) and are ecologically important in forest ecosystems as they contribute to wood decomposition and thus play essential roles in nutrient and carbon cycling (Seibold et al., 2021; Ulyshen, 2018). In addition, some species, such as the endangered great capricorn beetle (*Cerambyx cerdo*), act as keystone species within forest ecosystems, creating habitat for other species (Buse et al., 2008). In Europe, nearly 29.000 beetle species are recorded, including a large proportion of saproxylic beetles (Grove, 2002). One key driver for the high diversity of saproxylic insects is the diversity of tree species, including more than 80 tree genera in Europe (Mauri et al., 2017). Despite their ecological importance and close associations with host trees, the factors shaping saproxylic beetle diversity across tree taxa and larger geographical scales remain poorly understood. Existing studies suggest that beetle diversity can differ significantly between tree species and between native and non-native hosts (Gossner et al., 2016; Vogel et al., 2020). However, while some studies report host associations for saproxylic species, the degree and mechanisms of host specificity at the tree genus level remain debated (Milberg et al., 2014). Understanding how tree identity and associated traits influence saproxylic beetle diversity is therefore essential for explaining biodiversity patterns and for informing forest conservation and management.

Nevertheless, the robustness of these conclusions depends on the underlying evidence, much of which comes from literature-based datasets that may be affected by sampling bias. For example, Heilmann-Clausen et al. (2016) investigated drivers of wood-inhabiting fungal diversity, using both raw occurrence data and data standardized for sampling effort. While analyses of raw data supported previously reported patterns for saproxylic organisms (Brändle & Brandl, 2001; Strong et al., 1984), these relationships disappeared when controlling for sampling effort. This suggests that conclusions about ecological patterns may depend strongly on whether sampling effort is considered (Gotelli & Colwell, 2001). This is the case particularly in macroecological studies, where the use of species data from disparate areas and studies may result in the distortion of spatial patterns of biodiversity. A study conducted in China, for instance, demonstrated that inventory incompleteness has a significant impact on the explanatory power of environmental factors (Yang et al., 2013).

To better understand patterns of saproxylic beetle diversity, we quantified factors driving the diversity of saproxylic beetles across European tree genera and how distinct beetle communities are shaped among different hosts. Furthermore, we assess how our results are influenced by sampling effort and whether patterns derived from literature-based datasets differ from those obtained from standardized data. According to the *species-area hypothesis*, we predict that (P1) higher resource availability, reflected by greater host biomass, leads to increased saproxylic beetle diversity. We further expect (P2) larger trees to support higher beetle diversity, as they provide a greater number of microhabitats and microclimatic niches. Based on the *geological-time hypothesis*, we predict (P3) that tree genera with longer postglacial occurrence in Europe host a higher diversity of saproxylic beetles. Finally, following the *taxonomic-isolation hypothesis*, we expect (P4) tree genera with a higher number of congeneric species to support greater saproxylic beetle diversity.

## Materials and Methods

### Species data

We used the dataset from Kriegel et al. (2023) as our basis, expanded it, and supplemented it with five new datasets. New datasets were compiled from a range of experimental studies on saproxylic beetles conducted across Europe, encompassing a diverse array of forest types, from the Mediterranean to boreal forests (Figure 1). Only experimental setups that sampled for a sufficient duration to encompass the complete adult activity spectrum of all saproxylic species, and that allowed for the assignment of emerging saproxylic beetle species to a specific deadwood object, were considered.

**Figure 1:**
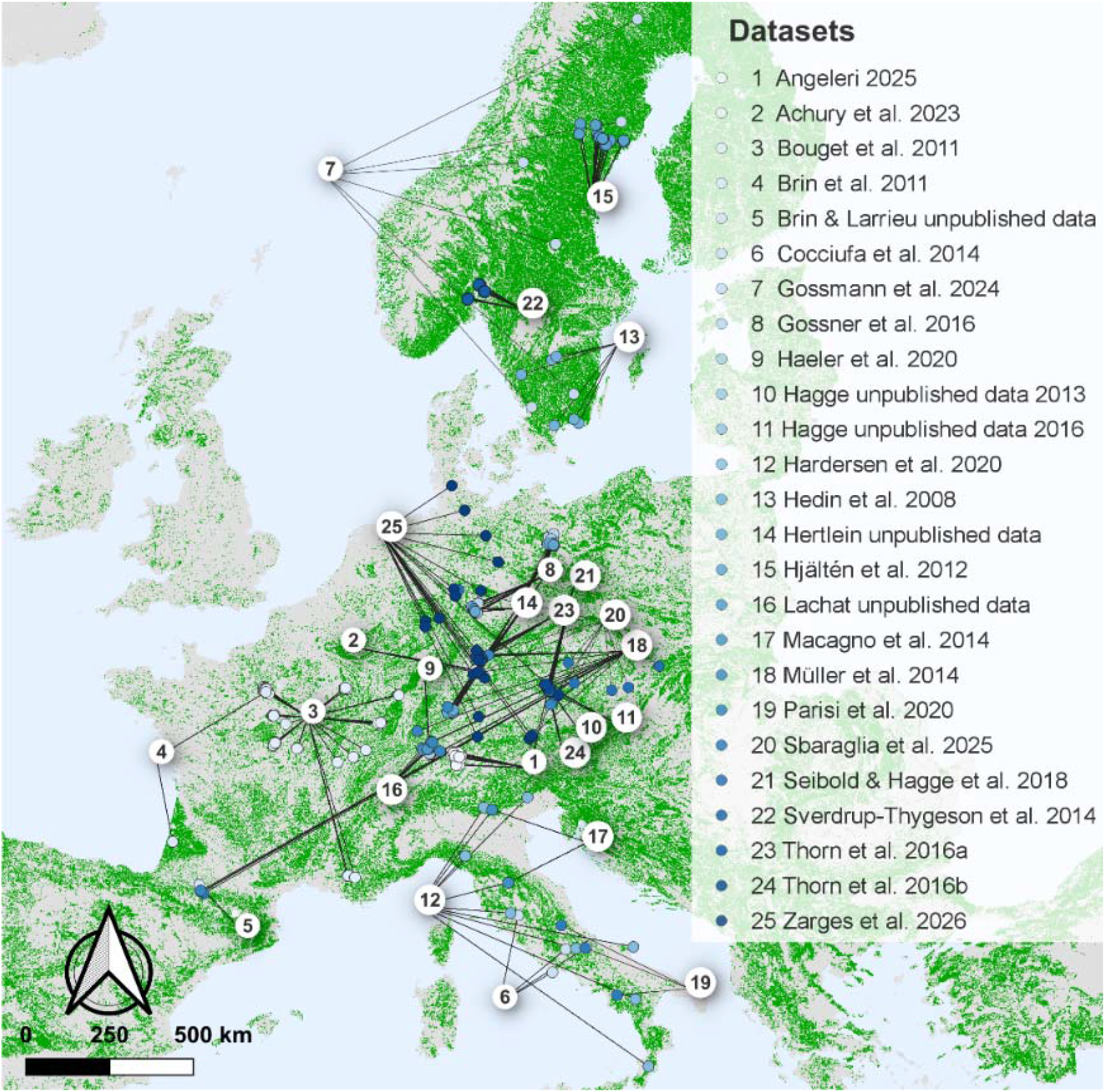
Distribution of the 25 datasets across Europe (List of dataset references are given in Supplementary Table). The background shows European forest cover taken from CORINE Land Cover 2018 (European Environment Agency, 2020).

**Figure 2:**
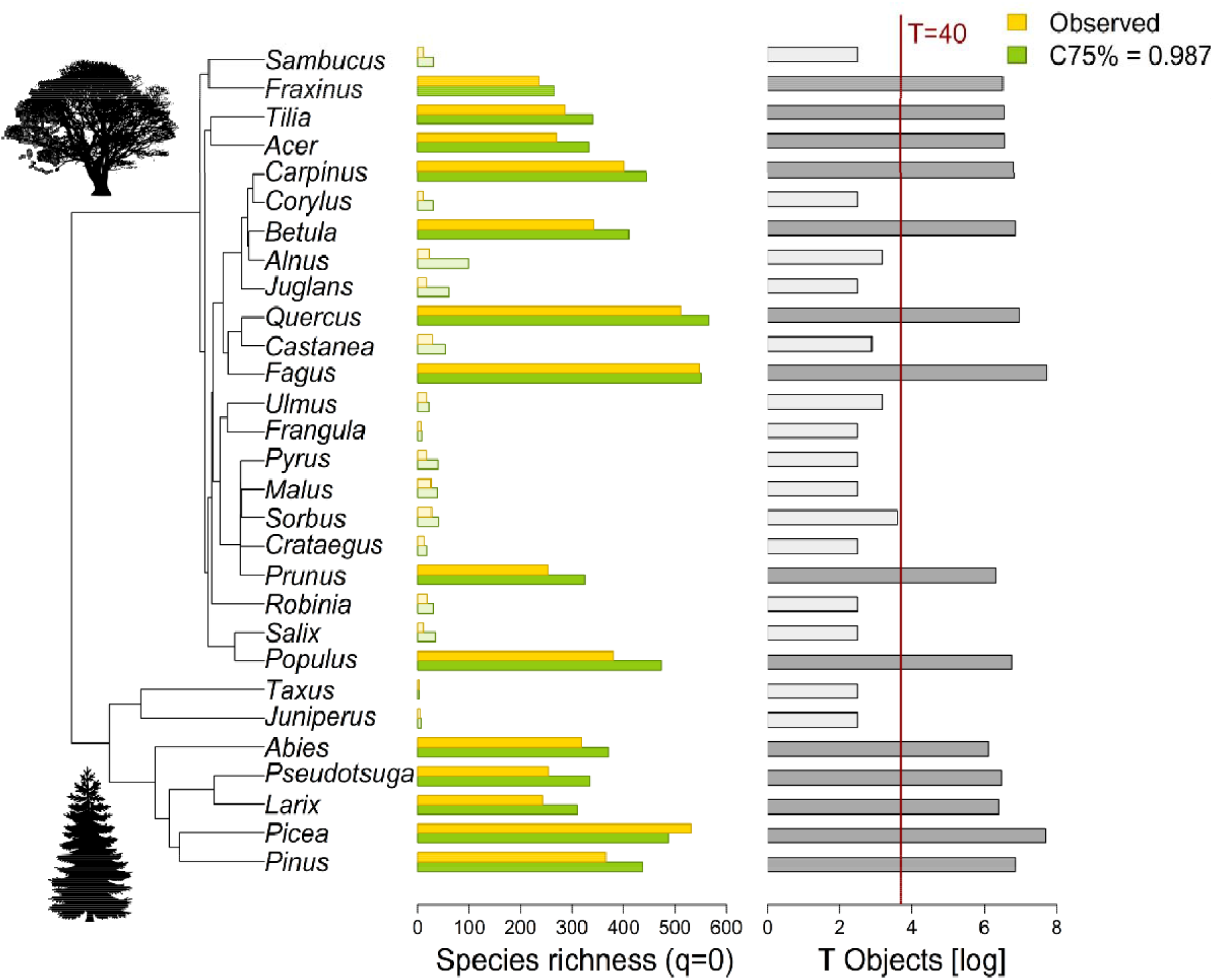
Coverage-based standardized beetle species richness (infrequent species, q = 0). The observed diversity is shown in yellow and the standardized beetle diversity for the coverage value C = 0.987 is shown in green. Grey bars represent the number of objects per tree genus, log transformed. Transparent bars represent all genera with less than 40 objects.

### Host tree traits

To identify potential main drivers of beetle species richness, we selected four variables: 1) the number of congeneric species as a measure of taxonomic isolation (Brändle & Brandl, 2001), 2) the host biomass as resource availability (basal area in m^2^) of each genus. Data from the German National Forest Inventory (Bwi, 2022) were used, as they provide a higher level of accuracy compared to a European-scale dataset (Mubareka et al., 2023), which is highly correlated (Supplementary Figure S1). 3) The mean maximum height of each genus was calculated using the maximum tree species height of each tree species provided by (Jäger et al., 2013) as a surrogate for structural complexity; and Johnson & More (2004). 4) The length of time of occurrence since the last glaciation in Europe (postglacial occurrence) (Brändle & Brandl, 2001) (Supplementary Table S2).

### Statistics

All statistical analyses were conducted in R version 4.4.1. Data were first pooled across study objects for each tree species. Given the absence of any knowledge regarding saproxylic beetle species relying exclusively on a single tree species, the data were further pooled at the genus level to improve the robustness of the analyses. To minimize the effect of intraspecific aggregation of beetle individuals within a deadwood object, each object was treated as a sampling unit, and only incidence (occurrence) records of beetle species were analyzed. Consequently, sample size corresponds to the number of deadwood objects, and the incidence frequency of each beetle species across objects was used as a proxy for its abundance. Accordingly, for incidence data, rare, common and dominant species are interpreted as infrequent, frequent and highly frequent species, respectively.

Because sampling completeness differed among tree genera, samples were standardized before comparing beetle diversity. To control for differences in sample completeness, we adopted the iNEXT (interpolation and extrapolation) framework (Chao et al., 2014), based on Hill numbers (Hill, 1973); see below), to standardize samples by sample coverage. Sample coverage is an objective measure of sample completeness and is defined as the proportion of all individuals (or occurrences) in the entire assemblage, including those of undetected species, that belong to the observed species.

Hill numbers provide a unified family of diversity measures that differ in their sensitivity to species relative abundances. Depending on the order of diversity (q), they emphasize different components of community structure. For q = 0, the Hill number equals species richness, treating all species equally regardless of their abundances. Species richness is particularly sensitive to the detection of rare or infrequent species and is therefore commonly regarded as a measure that emphasizes those. For q = 1, the Hill number (Shannon diversity) represents the effective number of common or frequent species. For q = 2, the Hill number (Simpson diversity) emphasizes dominant or highly frequent species.

Within the iNEXT framework, extrapolation can be extended only up to a maximum standardized coverage, denoted C-max, defined as the minimum estimated coverage at twice the observed sample size, SC(2T), across all samples, where *T* is the observed number of dead wood objects. Observed and standardized diversity for all Hill numbers were computed using the R package *iNEXT.3D* (Chao et al., 2021). When one or more samples exhibit very low coverage, *iNEXT.3D* instead uses 25th, 50th, or 75th percentiles of the SC(2T) values, denoted C25%, C50% and C75%, respectively, as the standardized coverage.

In our dataset of 29 tree genera (Supplementary Table S1), some genera were represented by relatively few deadwood objects, resulting in a low C-max value (close to 55%). After excluding genera with fewer than 40 objects (T < 40), the minimum SC(2T) is increased to approximately 98%. Consequently, all subsequent analyses were standardized to C75% = 0.987 using the subset of 14 tree genera (*Abies, Acer, Betula, Carpinus, Fagus, Fraxinus, Larix, Picea, Pinus, Populus, Prunus, Pseudotsuga, Quercus, Tilia*), each represented by at least 40 deadwood objects ().

To test whether literature-based species richness estimates are dependent on sampling effort, we compared the literature-based dataset of Müller et al. (2015) with our standardized saproxylic beetle species richness (q = 0). The correlation between the two datasets was assessed using Spearman’s rank correlation coefficient. To assess the potential influence of drivers, we selected saproxylic beetle diversity, along with Hill numbers, as the response variables and tree traits (conspecifics, resource availability, maximum tree height, and postglacial occurrence) as predictor variables. Before analyses, we examined the distribution of predictor variables (Supplementary Figure S11) and applied log transformation where necessary to reduce skewness and attain equal variance. However, host trees are not independent observation points due to their ancestral relationship. Because tree genera are not independent due to shared evolutionary history, we first tested for phylogenetic signal in species richness by estimating Pagel’s lambda (ranging from 0, indicating no constraint, to 1, indicating full constraint) based on the phylogeny provided by Durka & Michalski (2012) using the function phylosig in the package *phytools* (Revell, 2011). The estimated lambda was close to zero, indicating negligible phylogenetic signal. Consequently, subsequent analyses were performed using standard linear models with log-transformed diversity measures as the response. To identify the most important predictors, backward model selection based on Akaike’s Information Criterion (AIC) was applied using stepAIC from the *MASS* package (Ripley & Venables, 2009). Starting from the full model including all predictor variables, predictors were removed sequentially according to AIC. The model with the lowest AIC was retained. We extracted distance matrices based on fixed sample coverage by using the packages *iNEXT.beta3D* (Chao et al., 2023). We applied Nonmetric Multidimensional Scaling (NMDS) using the function metaMDS, to illustrate the community composition based on coverage-standardized dissimilarities and test those using PERMANOVA and pairwise R^2^ using the packages *vegan* (Oksanen et al., 2007) and *pairwiseAdonis* (Martinez Arbizu, 2020). Significance was assessed using 999 permutations. For calculating coverage-standardized dissimilarities the same number of sampling units is needed per genus. For each iteration, 435 objects per genus were randomly sampled without replacement. This procedure was repeated 5 times.

## Results

### Coverage-based standardized species richness

We compiled 25 datasets from 8 European countries covering 29 tree genera, 51 tree species, and 530,506 saproxylic beetles from 1041 species. In each study, singletons were present (except for the tree species *Taxus baccata*), indicating that data for all tree species are incomplete (Supplementary Material Table S1).

The observed species richness (q=0, infrequent species) is highest for *Fagus*, followed by *Picea*, followed by *Quercus*. Sample-size-based rarefaction and extrapolation curves for the diversity of infrequent species, exemplary for the most species-rich genera, revealed that sampling data did not contain sufficient information to accurately infer true diversity (Supplementary Material Figure S7). Standardizing the diversity of infrequent beetle species by coverage, the most species-rich genus was *Quercus*, followed by *Fagus*, when the coverage value was less than 99 % (). Comparing the standardized diversity of infrequent beetle species of our datasets from Germany with literature-based data, we found a correlation between both (Spearman rho = 0.701), with an underestimation of the genera *Fagus*, *Populus*, *Carpinus,* and *Pseudotsuga* in literature-based data, and an overestimation of *Pinus*, *Prunus*, *Larix,* and *Fraxinus* (Supplementary Material Figure S2).

The observed diversity of frequent beetle species (q = 1) was generally consistent with the pattern of the coverage-based standardized diversity of frequent beetle species (Sample-size-based rarefaction and extrapolation curves are shown for *Fagus, Quercus* and *Populus* in Supplementary Figure S7). This indicates that the data was sufficient to infer diversity of frequent species up to 100 %. The highest diversity of frequent beetle species was found for *Quercus,* followed by *Fagus* and *Populus* (Supplementary Figure S5).

The observed and coverage-based standardized diversity of highly frequent beetle species (q=2) showed the same pattern (Sample-size-based rarefaction and extrapolation curves are shown for *Fagus, Quercus,* and *Populus* in Supplementary Figure S6). The highest diversity of highly frequent beetle species was found for *Quercus,* followed by *Fagus* (Supplementary Figure S7).

### Drivers of saproxylic beetle diversity

Backward model selection based on AIC identified different sets of predictors along the Hill numbers. For observed infrequent beetle diversity, the final model included maximum tree height and postglacial occurrence, which showed negative effects, and resource availability, which showed positive effects on infrequent species diversity. However, for coverage-based standardized diversity of infrequent beetles, only maximum tree height and resource availability remained in the final model. For observed frequent beetle diversity, the final model retained only resource availability and showed a positive relationship with diversity. The same was found for standardized frequent beetle diversity. No predictor influenced observed and standardized highly frequent beetle diversity (Figure 3).

**Figure 3:**
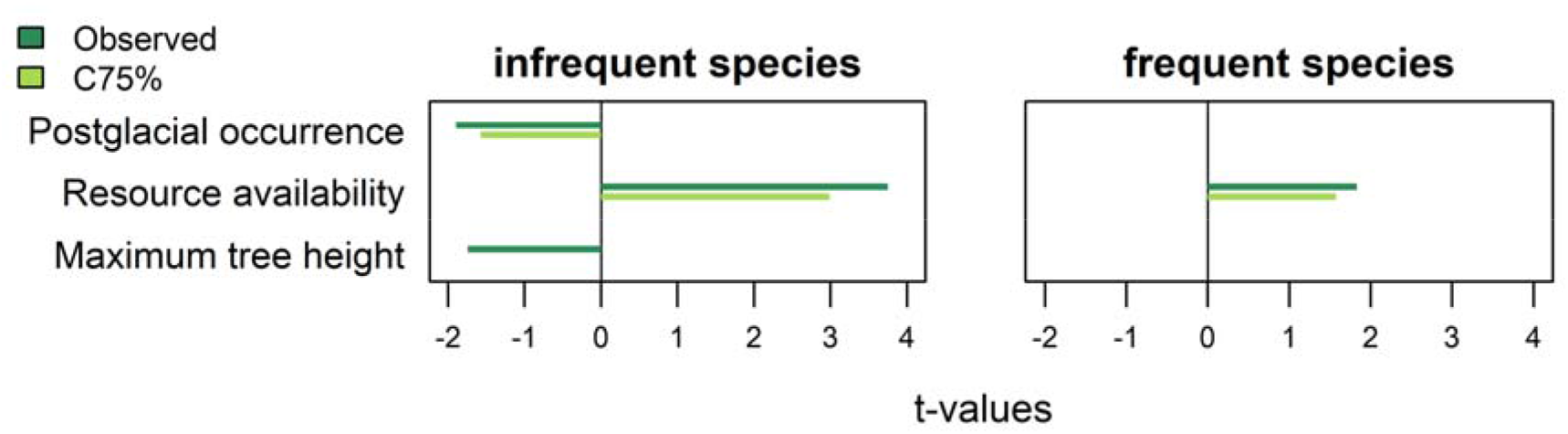
Linear model t-values of the predictors of the diversity of saproxylic beetle diversity along the Hill numbers for the observed coverage and for the coverage value C75 % = 0.987. Bars are shown for predictors selected by backward model selection based on AIC.

### Community composition of saproxylic beetles

Nonmetric Multidimensional Scaling was performed to compare the community composition of saproxylic beetles of 14 different tree genera. Using a subset of 435 objects per tree genus, observed data showed differences between tree genera similar for all, infrequent, frequent and highly frequent species, whereas standardization revealed clear compositional differences across Hill numbers (Figure 4, Supplementary Figure S8). Distinct community compositions were particularly apparent for *Abies*, *Fagus,* and *Picea*, whereas most other genera showed substantial overlap (Supplementary Figure S9). These visual patterns were supported by permutational multivariate analyses of variance, which detected a significant effect of tree genus on community composition across all Hill numbers. The strength of this effect increased from infrequent (R² = 0.39) to frequent (R² = 0.60) and highly frequent species (R² = 0.76). For infrequent saproxylic beetles, communities were mainly overlapping for most of the tree genera. Only *Abies* showed distinct assemblages across most comparisons in their beetle community composition. For frequent and highly frequent saproxylic beetle composition, separation increased markedly. Pairwise PERMANOVA revealed that *Abies, Fagus* and *Picea* supported clearly distinct communities with low within-genus variability, while *Quercus* exhibited comparatively high within-genus heterogeneity and overlapped with several other genera for infrequent community compositions but became more distinct when highly frequent species were considered (Figure 4, Supplementary Figure S9). Nonmetric Multidimensional Scaling was also performed for two other data subsets, which showed similar distinctions of communities between tree genera for infrequent, frequent and highly frequent species (Supplementary Figure S10).

**Figure 4:**
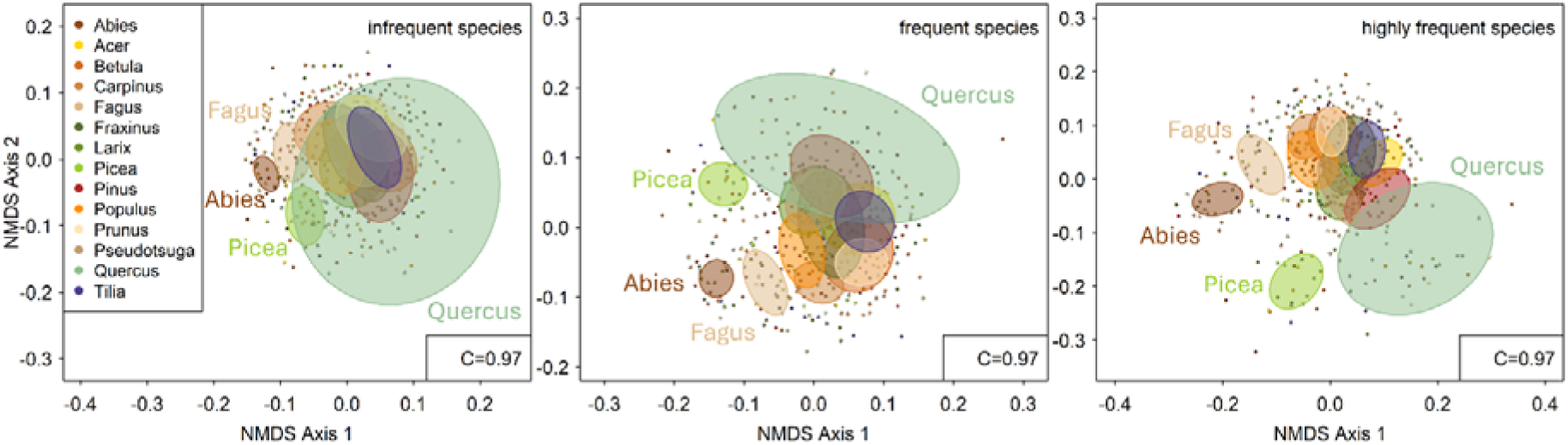
Differences in community compositions of saproxylic beetles along the Hill numbers, which can be interpreted as infrequent (q = 0), frequent (q = 1) and highly frequent (q = 2) species. Nonmetric Multidimensional Scaling (NMDS) was performed for a standardized coverage of C = 0.97. NMDS indicates the ordinal space within the first two axes of 435 objects, per tree genus.

Curculionidae made up the largest proportion of individuals overall, but their contribution varied strongly among tree genera. They accounted for over 60% in *Abies*, *Fagus*, and *Quercus*, and reached as high as 93% in *Picea*. In contrast, their proportion was much lower in *Populus* (10%), where Ciidae (32%) and Staphylinidae (13%) were more dominant.

In *Carpinus* and *Larix*, Curculionidae were only the second most abundant family, following Ciidae in *Carpinus* and Staphylinidae in *Larix*. While Staphylinidae were generally the second most abundant family across tree genera, an exception was *Quercus*, where Cerambycidae ranked second with 6%. Overall, the relative proportions of beetle families differed considerably between tree genera. (Figure 5, Supplementary Table S3).

**Figure 5:**
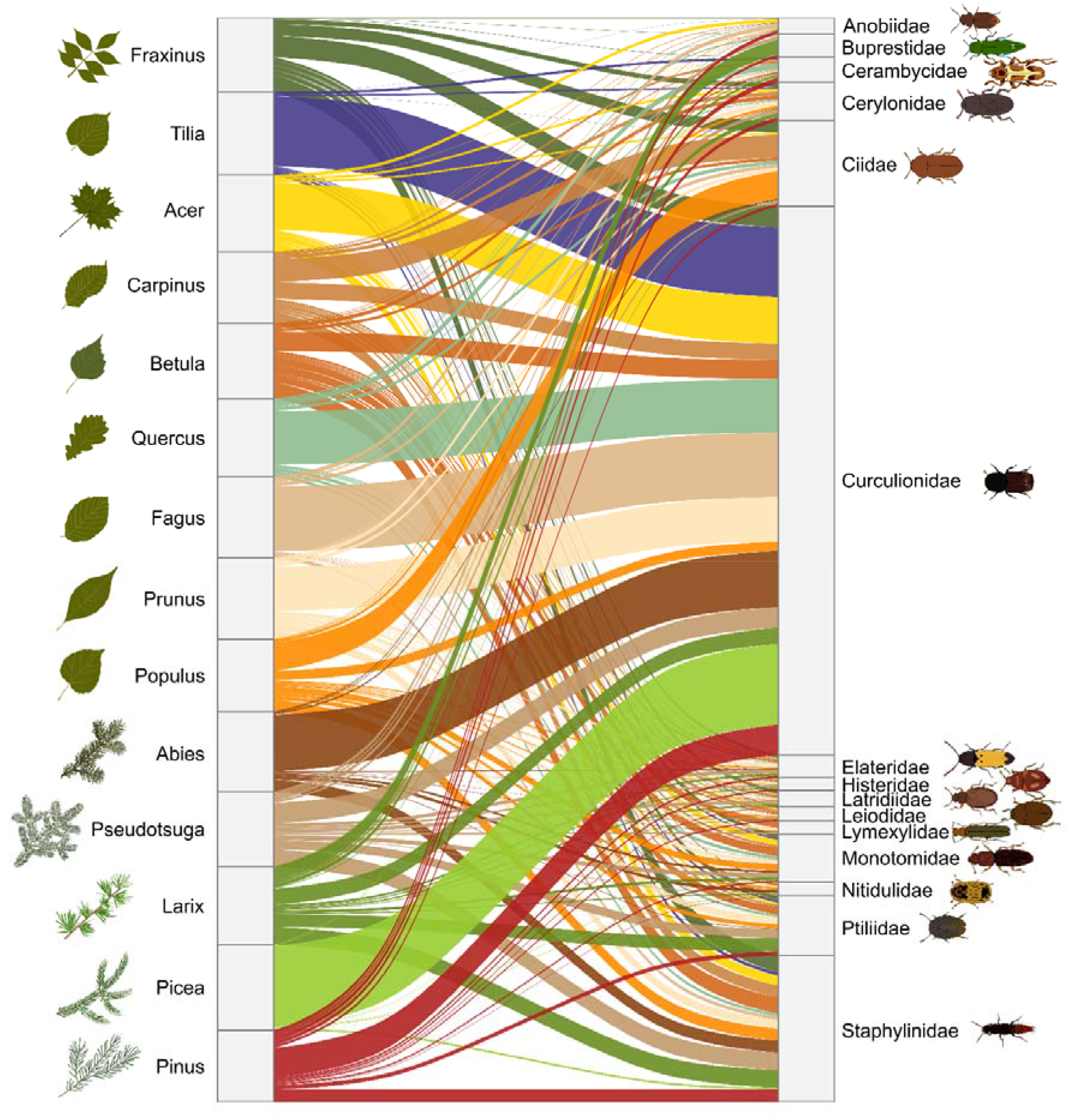
Community composition of the most abundant beetle families for 14 tree genera. The relative abundance of each beetle family was calculated for each tree genus, and the results are shown here for the 15 most abundant families.

## Discussion

Our study highlights *Quercus* as the most species rich European tree genus for saproxylic beetle diversity, supporting a disproportionately high diversity of infrequent species while also sustaining high diversity of frequent and highly frequent species. In contrast, the apparently high ranks of some other tree genera were partly explained by differences in sampling effort, emphasizing the importance of accounting for sampling bias when comparing host tree genera. Across genera, resource availability emerged as the primary driver of infrequent and frequent beetle diversity, whereas tree height influenced only infrequent species diversity. Contrary to expectations based on the *species-area hypothesis*, infrequent species diversity declined with increasing tree height. We further found only limited support for the *geological-time hypothesis*, after accounting for sampling completeness, the effect vanished. In addition, we found no evidence that taxonomic isolation of host trees shapes saproxylic beetle diversity. Finally, distinct beetle communities associated with several tree genera indicate that host tree identity contributes not only to species diversity but also to community composition.

### Differences in beetle diversity between tree genera

This study revealed *Quercus* to be the most species-rich tree genus for saproxylic beetles, followed by *Fagus* and *Picea* for infrequent species and by *Fagus* and *Populus* for frequent and highly frequent species. This finding is consistent with other studies that have identified oaks to have a high proportion of specialist and rare species compared to other tree genera (Jonsell et al., 1998; Müller & Gossner, 2007; Palm, 1959). Moreover, oak trees display not only a high level of species richness, but also a high proportion of monophagous species (Bussler, 2014; Jonsell et al., 1998; Milberg et al., 2014). Our NMDS revealed that saproxylic beetle assemblages overlap for infrequent species but become more distinct when highly frequent species were considered. This could be interpreted as meaning that infrequent species that occur on other genera can also be found on *Quercus* due to its broad habitat niches, while highly frequent species on oaks are more specialised and do not occur on other genera. One explanation for the high diversity of oaks could be that following the termination of the last glacial maximum, oaks demonstrated an exceptionally rapid recovery, due to more refugia during this time, establishing themselves much sooner than other species such as *Fagus sylvatica* (Firbas, 1949). But in our results, this effect vanished after standardising the data. *Quercus* is also distributed in the whole northern hemisphere, covering a wide ecological range from Mediterranean to hemiboreal areas (Nixon, 2006), which could lead to higher diversity based on the *species-area hypothesis* (Brändle & Brandl, 2001). But the same is true for the genera *Picea* (Yuan et al., 2025) and *Pinus* (Richardson, 2000). Another important factor could be the number of congeneric species. Within Europe, there are a similar number of species of *Quercus* (10) and *Pinus* (11) (Durka & Michalski, 2012), but our backward model selection did not identify congeneric species as explaining the diversity pattern of saproxylic beetles. However, focusing on a global scale, there are many more species within *Quercus* (Manos & Hipp, 2021) compared to *Pinus* (Richardson, 2000) and *Picea* (Yuan et al., 2025). Furthermore, oaks have a longer lifespan than other deciduous species, such as beech trees (Brown, 1996) and thus provide a stable habitat for insects. For instance, tree-related microhabitats (TreMs) – which promote specific species and/or communities (Larrieu et al., 2018) – such as rot-holes, are available for a longer period. And even if the tree is weakened, it will still remain standing for a longer period of time, despite the sanitary state of the bearing tree, which is assumed to decrease along its natural cycle (Larrieu et al., 2022).

*Fagus* had the second-highest coverage-based standardized beetle species richness. This is also in line with other studies, describing species of the genus *Fagus* with a saproxylic species diversity roughly as high as in *Quercus* (Walentowski et al., 2014). However, while a considerable number of saproxylic species colonize *Fagus*, only a small proportion exhibits a high degree of specialization towards this host tree. It has been estimated that approximately 65% of species may also occur in the absence of beech trees (Walentowski et al., 2010).

The highest diversity for frequent (q = 1) and highly frequent beetle (q = 2) species was found for *Quercus, Fagus and Populus. Populus* is considered a fast-growing, early-maturing and short-lived tree genus (Castillo et al., 2023; Stokland et al., 2012a). This makes it a suitable habitat for numerous saproxylic beetle species, especially pioneer species. Rapid decomposition quickly creates suitable TreMs (Larrieu et al., 2022) with varying degrees of decomposition on a single tree, allowing species with different habitat requirements to coexist on it. We also observed a high diversity of saproxylic beetles on *Carpinus*, characterized by a particularly high proportion of Ciidae. This pattern is likely driven by the rapid decomposition rate of *Carpinus* wood, which facilitates the early establishment of wood-decaying fungi (Kahl et al., 2017). As obligate fungivores, Ciidae rely on these fungal fruiting bodies and mycelia as essential microhabitats, explaining their increased abundance on this genus compared to slower-decaying substrates.

### Drivers of saproxylic beetle diversity on tree genera

Resource availability is frequently described as a major driver of saproxylic beetle diversity, with higher amounts of deadwood generally leading to increased species richness (Lassauce et al., 2011; Müller & Bütler, 2010; Seibold et al., 2016). Our results support this relationship, as increasing resource availability was associated with higher richness of infrequent and frequent saproxylic beetles. This finding is consistent with the *species-area hypothesis*, predicting that a larger amount of available habitat or resources supports more species (Brändle & Brandl, 2001). However, the *species-area hypothesis* also predicts that larger trees should host more species as they provide a greater number and diversity of microhabitats (Brändle & Brandl, 2001). In contrast to this expectation, our results show the opposite pattern. Increasing tree height was associated with a decrease in infrequent beetle species diversity. A similar pattern is reflected in the tree genera with the highest standardized beetle diversity in our dataset, such as *Quercus* and *Populus,* which tend to belong to relatively smaller tree genera (Jäger et al., 2013). This suggests that factors other than simple tree size, such as microhabitats, may be more important. Those are driven mainly by tree diameter and bark thickness, and therefore with tree age (Larrieu & Cabanettes, 2012; Vuidot et al., 2011), rather than tree height. For observed data, we additionally detected a negative relationship between postglacial occurrence and infrequent beetle diversity, meaning that tree genera with a shorter time since recolonisation hosted fewer infrequent beetle species, which supports the *geological-time hypothesis* (Brändle & Brandl, 2001). However, this effect disappeared after standardizing the data. This may indicate that the signal predicted by the *geological-time hypothesis* is partly driven by sampling bias rather than reflecting a true ecological mechanism. This highlights that apparent biogeographic relationships can be sensitive to differences in sampling completeness and that such differences need to be considered when interpreting large-scale patterns of host-associated biodiversity. Finally, we found no effect of the presence of congeneric tree species on saproxylic beetle diversity in either the observed or standardized datasets.

### Saproxylic beetle community composition for different tree genera

We found distinct community compositions for the tree genera *Abies, Fagus,* and *Picea.* With a high proportion of Curculionidae for most of the tree genera. Most saproxylic species tend to specialize on either conifers or broadleaves (Dahlberg & Stokland, 2004; Gossner et al., 2016; Müller et al., 2015; Stokland, 2012; Vogel et al., 2021). This may be attributed to different defensive mechanisms in coniferous and deciduous trees. One reason could be the differences in anatomical and physiochemical properties of deciduous and coniferous tree species (Meerts, 2002; Weedon et al., 2009). For instance, the resin defense of conifers seals wounds within a few hours, whereas those of deciduous trees may remain exposed for longer periods, providing colonisation opportunities to a diverse array of wound-associated species (Stokland et al., 2012b). Moreover, coniferous and broadleaved trees support distinct fungal communities (Kebli et al., 2011; Van Der Wal et al., 2016). Given that many saproxylic beetles are fungivorous and closely linked to these fungi (Jonsell et al., 1998; Kirisits, 2007), this specialization may contribute further to these differences. At the level of narrower specializations, saproxylic species demonstrate a range of degrees of host specificity (Bussler et al., 2011; Speight, 1989). Ehnström & Axelsson (2002) showed that a large proportion of bark beetles (Scolytinae, Curculionidae) are specialized on tree genera, which could explain the distinct communities for those genera. We also found that *Fagus* and *Picea* are part of the most diverse genera for infrequent saproxylic beetle species, which could also lead to the distinct community compositions. However, we found assemblages of infrequent species largely overlapping across all tree genera. This could indicate that infrequent species are not dependent on the tree species, but on rare structures, such as late decay stages or rare TreM-types. TreMs in particular represent essential substrates for a wide range of taxa, including saproxylic insects and fungi, and can drive species occurrence independently of tree species identity (Larrieu et al., 2018). Thus, tree genus appears to be important for structuring saproxylic beetle communities, but its importance differs among components of biodiversity. Host identity may be particularly relevant for frequent species, whereas infrequent species may depend more strongly on habitat structures that occur across multiple tree genera.

### Conservation implications

Our results show that tree genus is an important determinant of saproxylic beetle diversity at the European scale. *Quercus* supported the highest standardized richness of infrequent, frequent and highly frequent species, while *Picea, Fagus* and *Abies* supported a distinct community despite their high diversity. Maintaining a diversity of tree genera may therefore support both high species richness and complementary beetle communities. The contribution of tree genera also depended on their availability across forest landscapes. Resource availability was the strongest predictor of beetle diversity. Conservation should therefore consider both the diversity and availability of tree genera rather than focusing on individual genera alone. Finally, sampling completeness strongly affected observed diversity patterns. The relationship between postglacial occurrence and infrequent beetle diversity disappeared after accounting for sampling completeness, indicating that apparent biogeographic patterns can partly reflect differences in sampling effort. Coverage-based standardization should therefore be considered when comparing saproxylic beetle diversity among tree genera.

## Conclusion

This study demonstrates that tree genus is a key determinant of saproxylic beetle diversity across Europe and further show that standardization of sampling effort substantially alters observed diversity patterns. Thus, the *geological-time hypothesis* may reflect sampling effects rather than true ecological patterns. *Quercus* and *Fagus* supported the highest richness of infrequent, frequent and highly frequent species, indicating a wide range of ecological niches for saproxylic beetles. Only a limited number of distinct beetle communities were associated with coniferous trees. Nevertheless, *Picea* harboured a distinct community while maintaining high diversity, highlighting its complementary role for conserving specialized species. Overall, individual trees can host a broad variety of beetle assemblages, many of which also occur on other tree genera. From a conservation perspective, maintaining a diverse composition of tree genera remains important because tree identity stringly influeces saproxylic beetle diversity and community composition.However, tree genera alone are not sufficient, as they do not necessarily promote beetle communities of infrequent species; for these, the focus should instead be placed on microhabitat diversity. In addition, standardized data revealed higher richness on smaller trees, contradicting its prediction. Instead of tree size, the occurrence of different niches, due to microhabitats, may be more important. Forest management strategies should therefore prioritize the retention and regeneration of diverse, microhabitat-rich stands, with special consideration for genera such as *Quercus*, *Fagus* and *Picea,* which support particularly high or complementary levels of diversity. Such approaches are expected to maximize saproxylic beetle diversity at the stand and landscape scale. However, some tree genera are likely underrepresented in current datasets, potentially obscuring important host-diversity relationships. Targeted sampling of these underrepresented genera is therefore needed to obtain a more complete understanding of drivers of saproxylic beetle diversity.

## Acknowledgements

The study received funding from the DBU (Deutsche Bundesstiftung Umwelt). The unpublished dataset from M. Hertlein has been funded by the DFG Priority Program 1374 Biodiversity-Exploratories (project number 512286464).

