## Supplement for "Sample coverage standardization ranks species richness of saproxylic beetles across European tree genera"

*Supplementary Material*

Supplement Table S1: **Data summary for beetle species on 29 tree genera based on 25 studies**. n = total abundance (total number of beetles); S.obs = observed number of species; qTD = estimated species richness (q=0) for a sample coverage of 0.987; SC(T) = sample coverage for the reference sample of size T; SC(2T) = sample coverage for double the reference sample size; Q1 = number of uniques (i.e., species detected in only one object); f2 = number of duplicates (i.e., species detected in two objects); T: number of objects; number of datasets.

| Tree genus | n | S.obs | qTD C75% | SC(T) | SC(2T) | Q1 | Q2 | T (number of objects) | Number of Datasets |
| --- | --- | --- | --- | --- | --- | --- | --- | --- | --- |
| *Abies* | 21490 | 319 | 369.94 | 0.969 | 0.989 | 92 | 47 | 449 | 6 |
| *Acer* | 7370 | 271 | 333.10 | 0.955 | 0.984 | 91 | 47 | 714 | 3 |
| *Alnus* | 369 | 24 | 99.29 | 0.565 | 0.654 | 18 | 2 | 24 | 1 |
| *Betula* | 6464 | 342 | 410.28 | 0.969 | 0.986 | 102 | 44 | 958 | 5 |
| *Carpinus* | 15550 | 403 | 446.05 | 0.977 | 0.991 | 103 | 52 | 927 | 4 |
| *Castanea* | 114 | 28 | 54.28 | 0.494 | 0.755 | 20 | 7 | 18 | 2 |
| *Corylus* | 77 | 11 | 29.17 | 0.765 | 0.836 | 7 | 0 | 12 | 1 |
| *Crataegus* | 65 | 13 | 18.34 | 0.724 | 0.915 | 7 | 4 | 12 | 1 |
| *Fagus* | 113367 | 547 | 550.37 | 0.986 | 0.994 | 128 | 62 | 2301 | 17 |
| *Frangula* | 39 | 7 | 8.26 | 0.841 | 0.978 | 3 | 3 | 12 | 1 |
| *Fraxinus* | 3301 | 237 | 265.34 | 0.963 | 0.991 | 64 | 46 | 674 | 2 |
| *Juglans* | 45 | 18 | 61.84 | 0.454 | 0.598 | 14 | 2 | 12 | 1 |
| *Juniperus* | 16 | 5 | 6.70 | 0.816 | 0.935 | 2 | 1 | 12 | 1 |
| *Larix* | 3445 | 242 | 311.00 | 0.944 | 0.978 | 86 | 41 | 604 | 2 |
| *Malus* | 2370 | 26 | 38.27 | 0.814 | 0.922 | 12 | 5 | 12 | 1 |
| *Picea* | 265460 | 532 | 488.12 | 0.990 | 0.995 | 135 | 58 | 2220 | 13 |
| *Pinus* | 7805 | 366 | 437.52 | 0.963 | 0.986 | 107 | 51 | 956 | 6 |
| *Populus* | 11053 | 380 | 473.47 | 0.965 | 0.983 | 115 | 44 | 858 | 4 |
| *Prunus* | 6879 | 253 | 324.94 | 0.951 | 0.980 | 90 | 41 | 555 | 2 |
| *Pseudotsuga* | 3343 | 254 | 334.23 | 0.948 | 0.977 | 88 | 36 | 657 | 2 |
| *Pyrus* | 259 | 18 | 39.32 | 0.574 | 0.750 | 12 | 3 | 12 | 1 |
| *Quercus* | 39159 | 511 | 566.28 | 0.974 | 0.992 | 139 | 84 | 1068 | 9 |
| *Robinia* | 264 | 19 | 29.62 | 0.692 | 0.881 | 11 | 5 | 12 | 1 |
| *Salix* | 17 | 12 | 34.48 | 0.310 | 0.551 | 10 | 2 | 12 | 1 |
| *Sambucus* | 54 | 12 | 30.08 | 0.491 | 0.683 | 9 | 2 | 12 | 1 |
| *Sorbus* | 306 | 27 | 40.17 | 0.874 | 0.940 | 11 | 4 | 36 | 1 |
| *Taxus* | 19 | 3 | 3.20 | 0.853 | 0.996 | 1 | 2 | 12 | 1 |
| *Tilia* | 21521 | 287 | 341.14 | 0.960 | 0.986 | 88 | 48 | 707 | 3 |
| *Ulmus* | 285 | 18 | 22.72 | 0.825 | 0.961 | 8 | 6 | 24 | 1 |

Supplement Table 2: Host tree traits. Values of the four potential drivers of saproxylic beetle species diversity for a subset of 14 tree genera.

| **Tree genus** | **Resource [m^2^]** | **Postglacial occurrence [log]** | **# congeneric species** | **Maximum tree height** |
| --- | --- | --- | --- | --- |
| ***Abies*** | 81828006.4 | 5000 | 4 | 60 |
| ***Acer*** | 40255663.3 | 8000 | 6 | 23.33 |
| ***Betula*** | 69510428.7 | 14000 | 5 | 25 |
| ***Carpinus*** | 31676558 | 4000 | 1 | 20 |
| ***Fagus*** | 583355957 | 6000 | 1 | 40 |
| ***Fraxinus*** | 58709994.8 | 8000 | 3 | 40 |
| ***Larix*** | 91425934 | 11000 | 3 | 35 |
| ***Picea*** | 1231108099 | 5200 | 5 | 50 |
| ***Pinus*** | 704839512 | 14000 | 11 | 40 |
| ***Populus*** | 31710588.7 | 12000 | 8 | 25 |
| ***Prunus*** | 8867109.53 | 12000 | 18 | 13.33 |
| ***Pseudotsuga*** | - | - | 1 | 90 |
| ***Quercus*** | 302378159 | 10000 | 10 | 31 |
| ***Tilia*** | 10736884.9 | 9000 | 4 | 25 |

Supplement Table S3: **Abundance of saproxylic beetles per tree genus**. Abundance of the 15 most abundant beetle families.

|  |  | *Abies* | *Acer* | *Betula* | *Carpinus* | *Fagus* | *Fraxinus* | *Larix* | *Picea* | *Pinus* | *Populus* | *Prunus* | *Pseudotsuga* | *Quercus* | *Tilia* |
| --- | --- | --- | --- | --- | --- | --- | --- | --- | --- | --- | --- | --- | --- | --- | --- |
| Abundance of beetle families | **Anobiidae** | 29 | 275 | 35 | 111 | 5552 | 31 | 21 | 72 | 335 | 87 | 24 | 15 | 319 | 29 |
|  | **Bupresti-dae** | 4 | 0 | 11 | 92 | 170 | 11 | 576 | 768 | 227 | 3 | 2 | 133 | 594 | 1 |
|  | **Ceramby-cidae** | 259 | 28 | 67 | 425 | 1884 | 16 | 86 | 1549 | 443 | 77 | 43 | 76 | 2590 | 625 |
|  | **Ceryloni-dae** | 184 | 209 | 299 | 665 | 633 | 187 | 167 | 365 | 345 | 336 | 372 | 139 | 563 | 392 |
|  | **Ciidae** | 357 | 161 | 221 | 4141 | 4609 | 443 | 31 | 220 | 230 | 3566 | 267 | 87 | 1577 | 39 |
|  | **Curculio-nidae** | 13968 | 3974 | 1404 | 2927 | 84475 | 809 | 637 | 242392 | 2698 | 1196 | 3499 | 789 | 24232 | 17273 |
|  | **Elateridae** | 285 | 65 | 154 | 363 | 445 | 96 | 86 | 393 | 176 | 181 | 260 | 130 | 421 | 185 |
|  | **Histeridae** | 43 | 46 | 104 | 246 | 172 | 27 | 44 | 917 | 147 | 108 | 83 | 74 | 513 | 72 |
|  | **Latridiidae** | 38 | 61 | 131 | 205 | 361 | 139 | 59 | 295 | 69 | 114 | 68 | 127 | 375 | 66 |
|  | **Leiodidae** | 68 | 161 | 111 | 609 | 181 | 44 | 28 | 78 | 124 | 174 | 71 | 33 | 122 | 114 |
|  | **Lymexylidae** | 130 | 4 | 519 | 6 | 607 | 5 | 1 | 11 | 2 | 100 | 178 | 82 | 6 | 3 |
|  | **Monoto-midae** | 535 | 478 | 621 | 434 | 1262 | 156 | 109 | 1259 | 137 | 835 | 280 | 169 | 1508 | 414 |
|  | **Nitidulidae** | 794 | 48 | 159 | 60 | 511 | 34 | 49 | 1619 | 39 | 343 | 47 | 27 | 65 | 43 |
|  | **Ptiliidae** | 164 | 191 | 552 | 380 | 441 | 170 | 507 | 392 | 382 | 543 | 554 | 342 | 1184 | 547 |
|  | **Staphylini-dae** | 2944 | 831 | 1203 | 2116 | 4769 | 643 | 700 | 5638 | 1125 | 1520 | 688 | 673 | 1362 | 844 |

Supplement Table S4: **List of Dataset references used for analysis.**

Datasets

Angeleri et al. (2026)

Achury et al. (2023)

Bouget et al. (2011)

Brin et al. (2011)

Brin & Larrieu unpublished data

(Cocciufa et al., 2014)

(Goßmann et al., 2024)

(Gossner et al., 2016)

Haeler unpublished data 2013

Haeler unpublished data 2016

(Hardersen et al., 2020)

(Hedin et al., 2008)

Hertlein unpublished data

(Hjältén et al., 2012)

Lachat unpublished data

(Macagno et al., 2015)

(Müller et al., 2014)

(Parisi et al., 2020)

(Sbaraglia et al., 2026)

(Seibold et al., 2018)

(Sverdrup-Thygeson et al., 2014)

(Thorn et al., 2016a)

(Thorn et al., 2016b)

(Zarges et al., 2026)

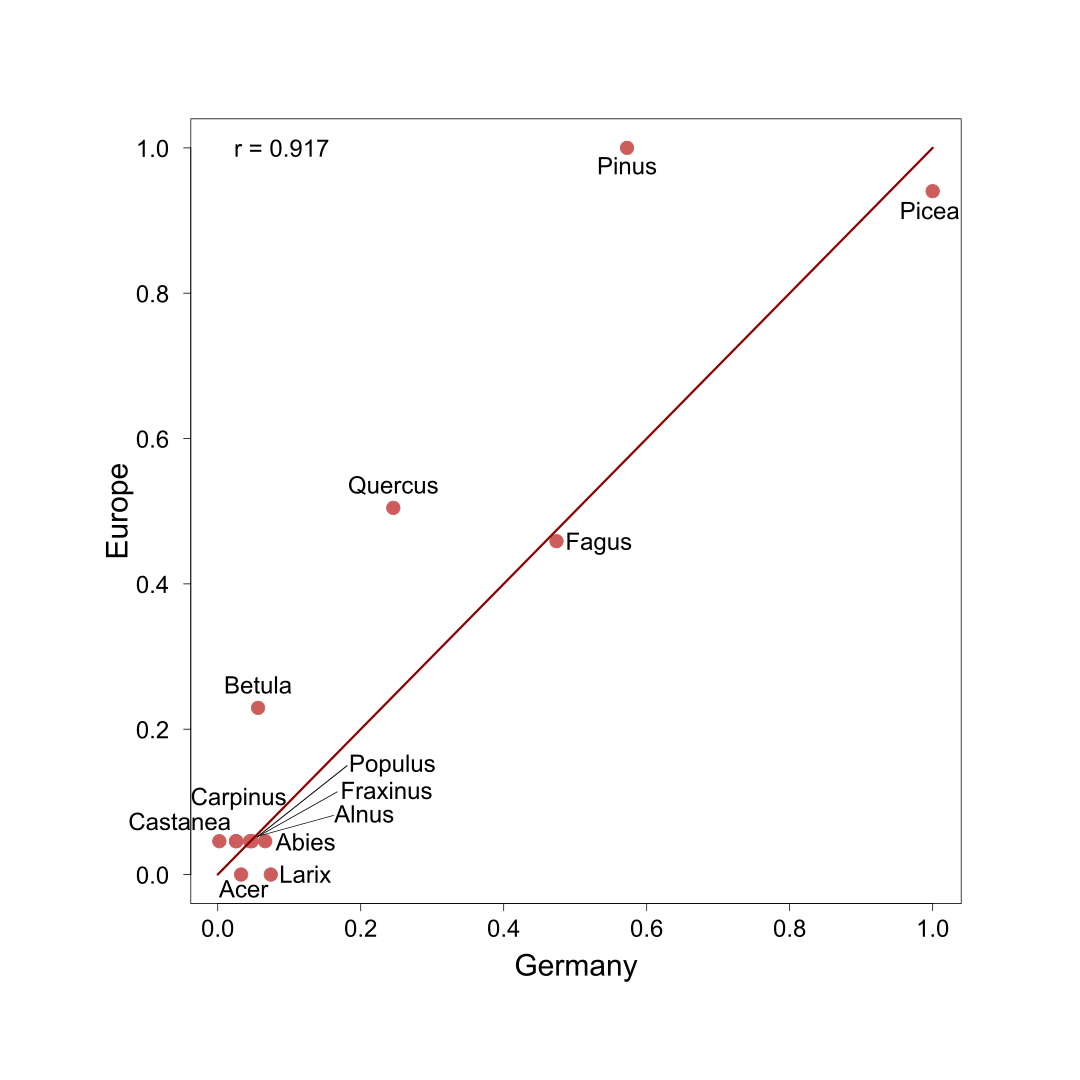

Supplement Figure S1: **Correlation between biomass datasets** of tree genera for Europe and Germany. For Europe the dataset Mubareka et al. (2023) and for Germany the BWI ([www.bundeswaldinventur.de](http://www.bundeswaldinventur.de)) dataset were used and a Pearson correlation was calculated.

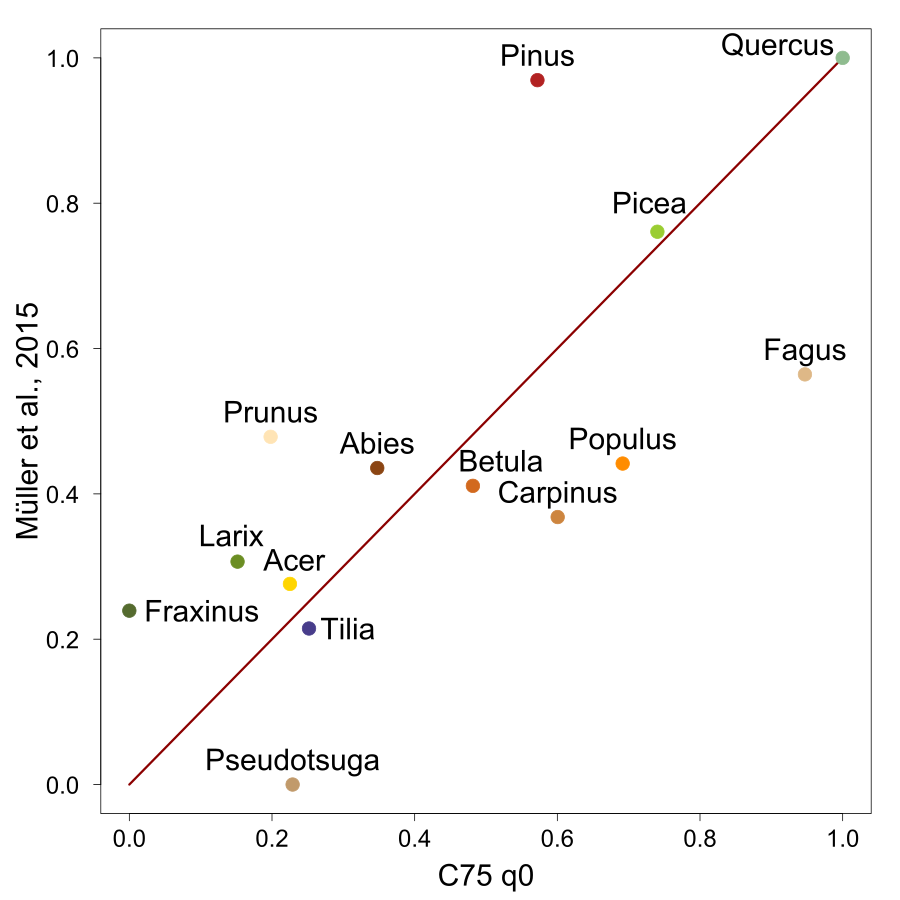

Supplement Figure S2: Correlation between literature-based (Müller et al., 2015) and standardized rare beetle diversity for a coverage of C75=0.987. The diversity ranks of rare saproxylic beetle species of Germany for a subset of 14 tree genera are shown. Genera above the red line are overestimated and those below are underestimated in the literature-based dataset

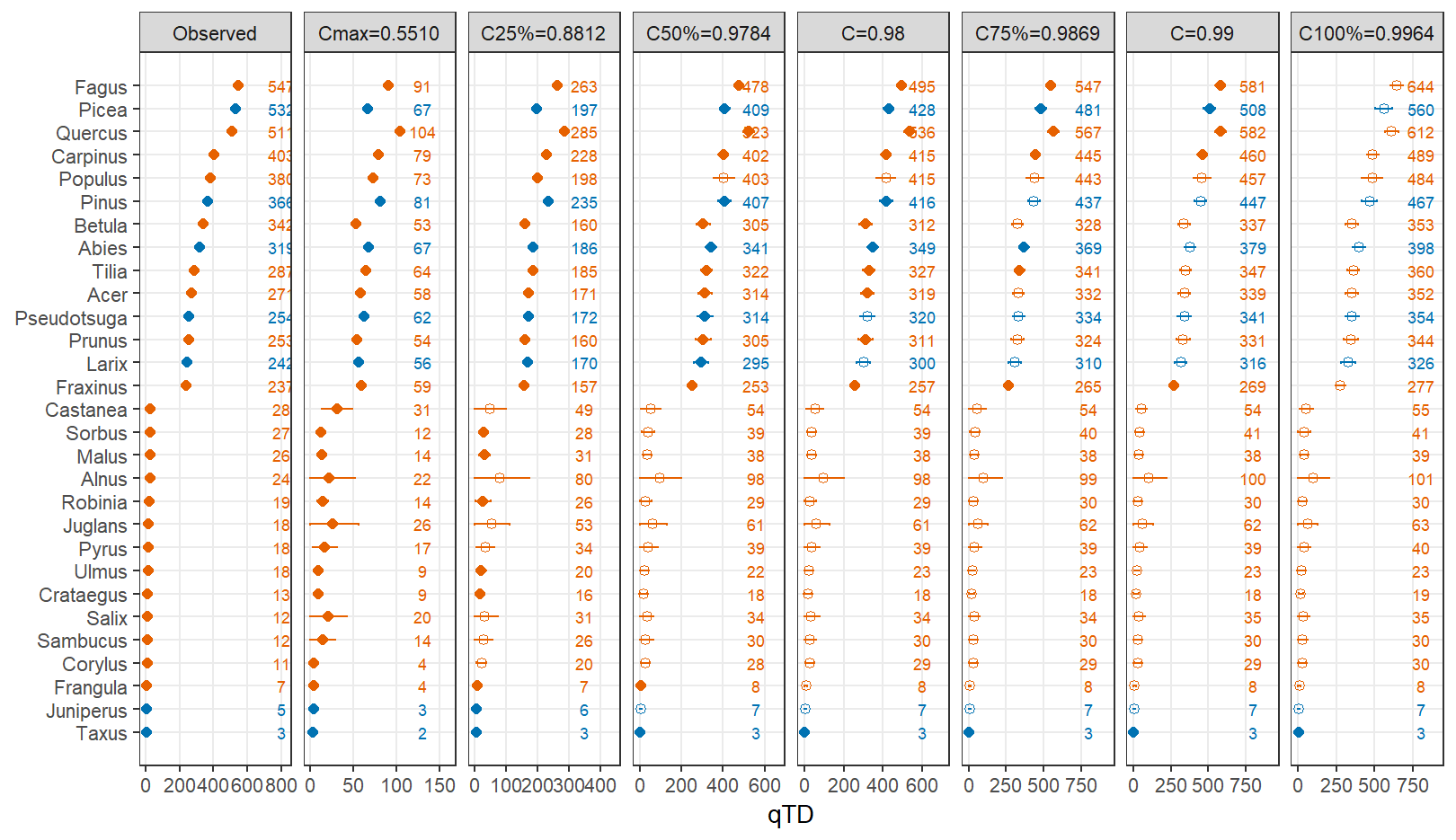
Supplement Figure S3: **(q = 0)** **4 Datasets of northern Europe excluded. Coverage-based standardized beetle species richness (numerical value on the right of each panel) with 95% confidence intervals (some are short so invisible) for several coverage values.** Tree genera are ordered by the observed species richness (Panel 1). Orange dots mean broadleaf tree genus; blue dots mean coniferous tree genus. Solid dots represent reliable extrapolation estimates because each extrapolated size is less than twice the observed sample size; hollow dots represent that extrapolated sample sizes exceed twice the observed sample sizes and thus may unreliable.

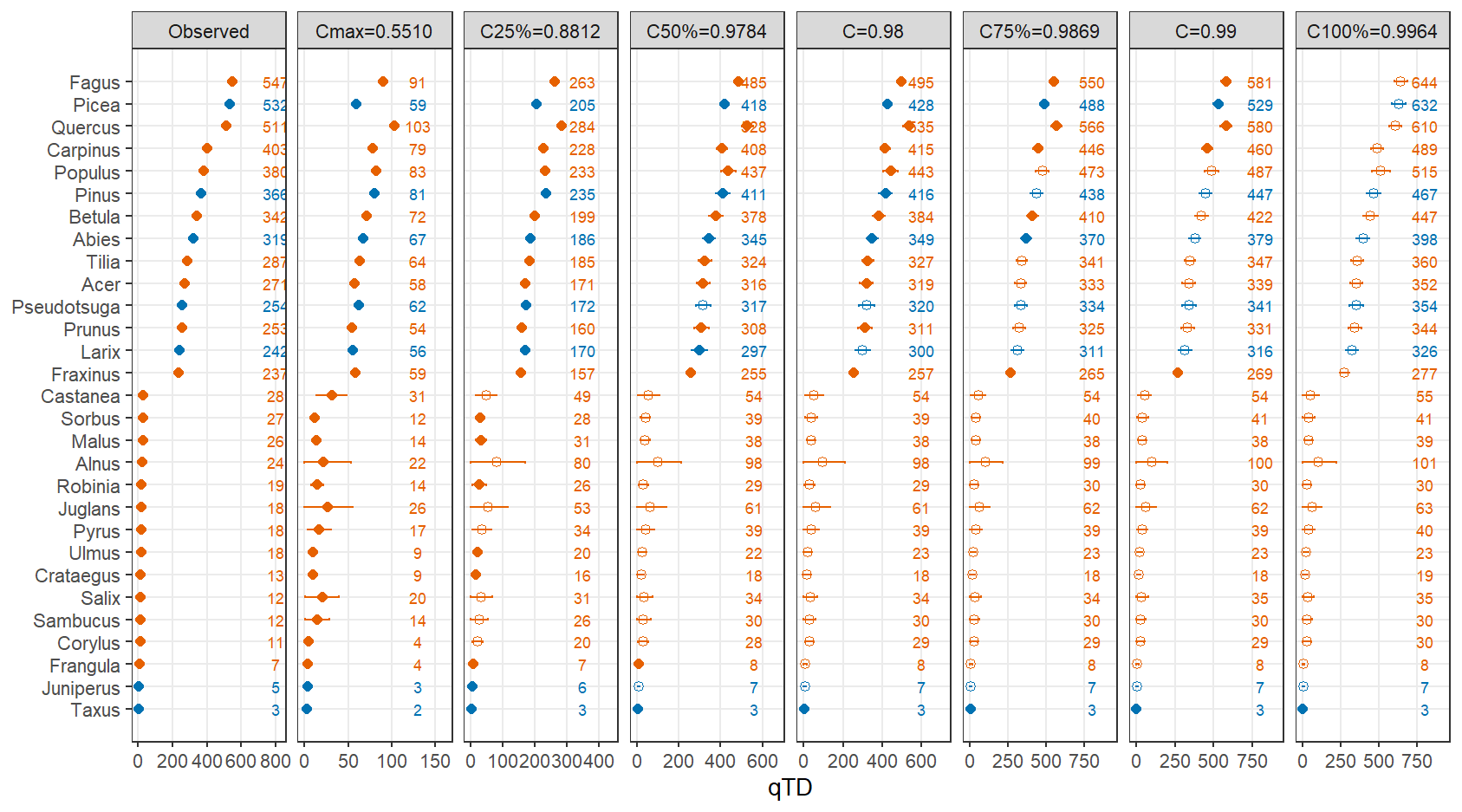
Supplement Figure S4: **(q = 0)** **Coverage-based standardized beetle species richness (numerical value on the right of each panel) with 95% confidence intervals (some are short so invisible) for several coverage values.** Tree species are ordered by the observed species genera (Panel 1). Orange dots mean broadleaf tree genus; blue dots mean coniferous tree genus. Solid dots represent reliable extrapolation estimates because each extrapolated size is less than twice the observed sample size; hollow dots represent that extrapolated sample sizes exceed twice the observed sample sizes and thus may be unreliable.

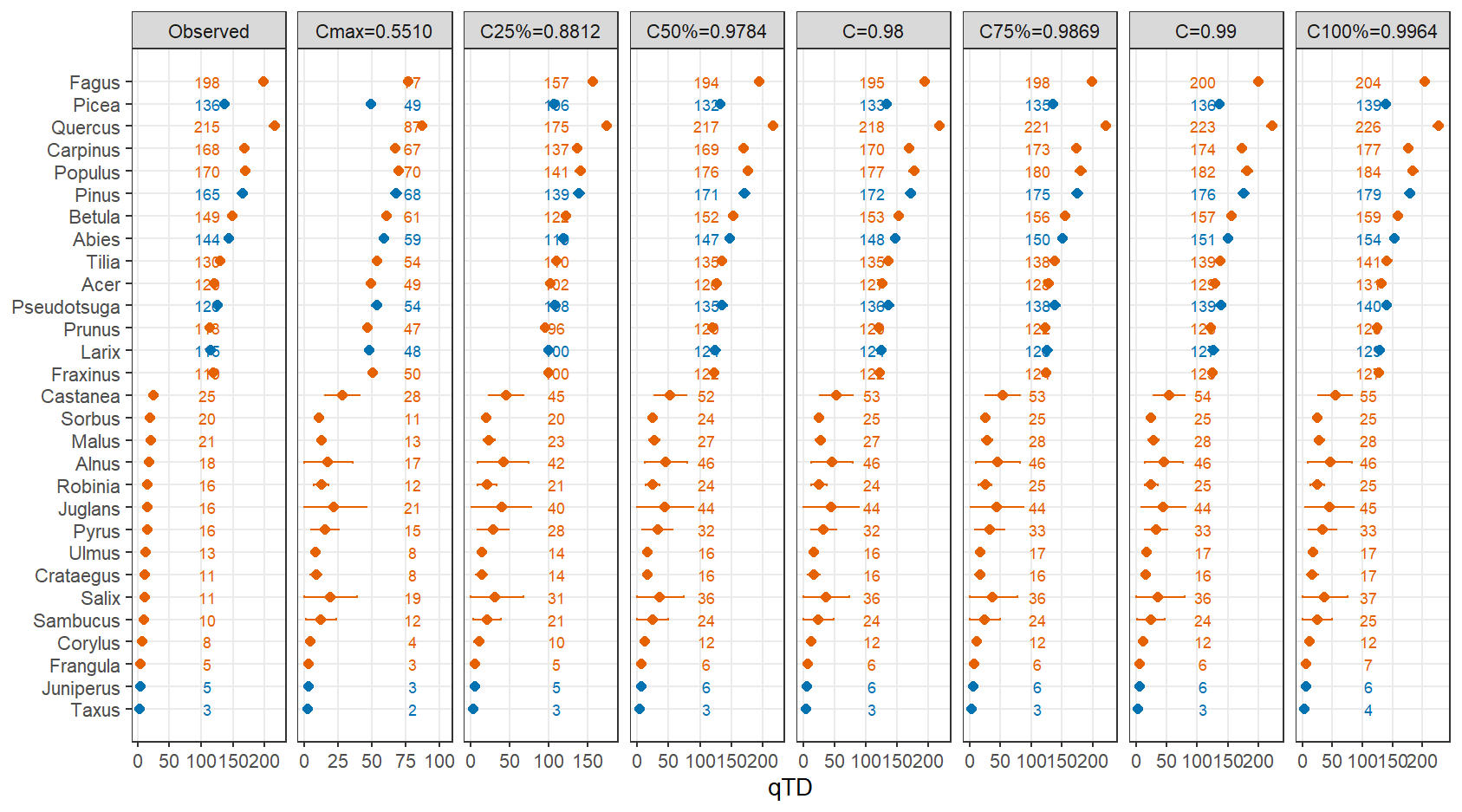
Supplement Figure S5: **(q = 1)** **Coverage-based standardized beetle Shannon diversity estimate (numerical value on the right of each panel) with 95% confidence intervals (some are short so intervals are invisible) for several coverage values.** Tree genera are ordered by the observed species richness (Panel 1 in Figure 2). Orange dots mean broadleaf tree genus; blue dots mean coniferous tree genus. For each coverage value, the standardized diversity pattern is generally consistent with the observed pattern, signifying that data are sufficient to infer diversity of q = 1 up to 100% (asymptotes).

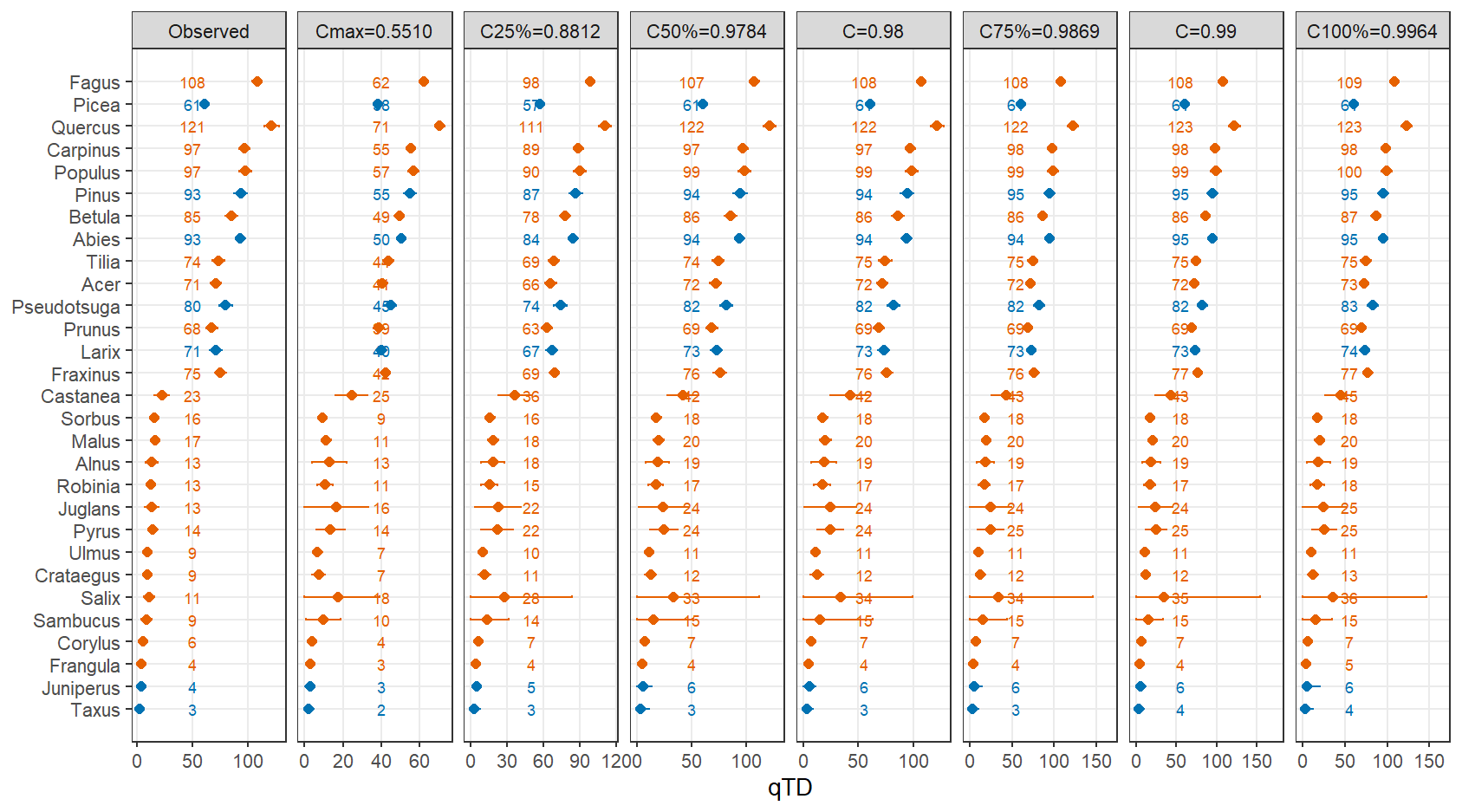

Supplement Figure S6: **(q = 2)** **Coverage-based standardized beetle Simpson diversity estimate (numerical value on the right of each panel) with 95% confidence intervals (some are short so intervals are invisible) for several coverage values.** Tree genera are ordered by the observed species richness (Panel 1 in Figure 2). Orange dots mean broadleaf tree genus; blue dots mean coniferous tree genus. For each coverage value, the standardized diversity pattern is generally consistent with the observed pattern, signifying that data are sufficient to infer diversity of q = 2 up to 100% (asymptotes).

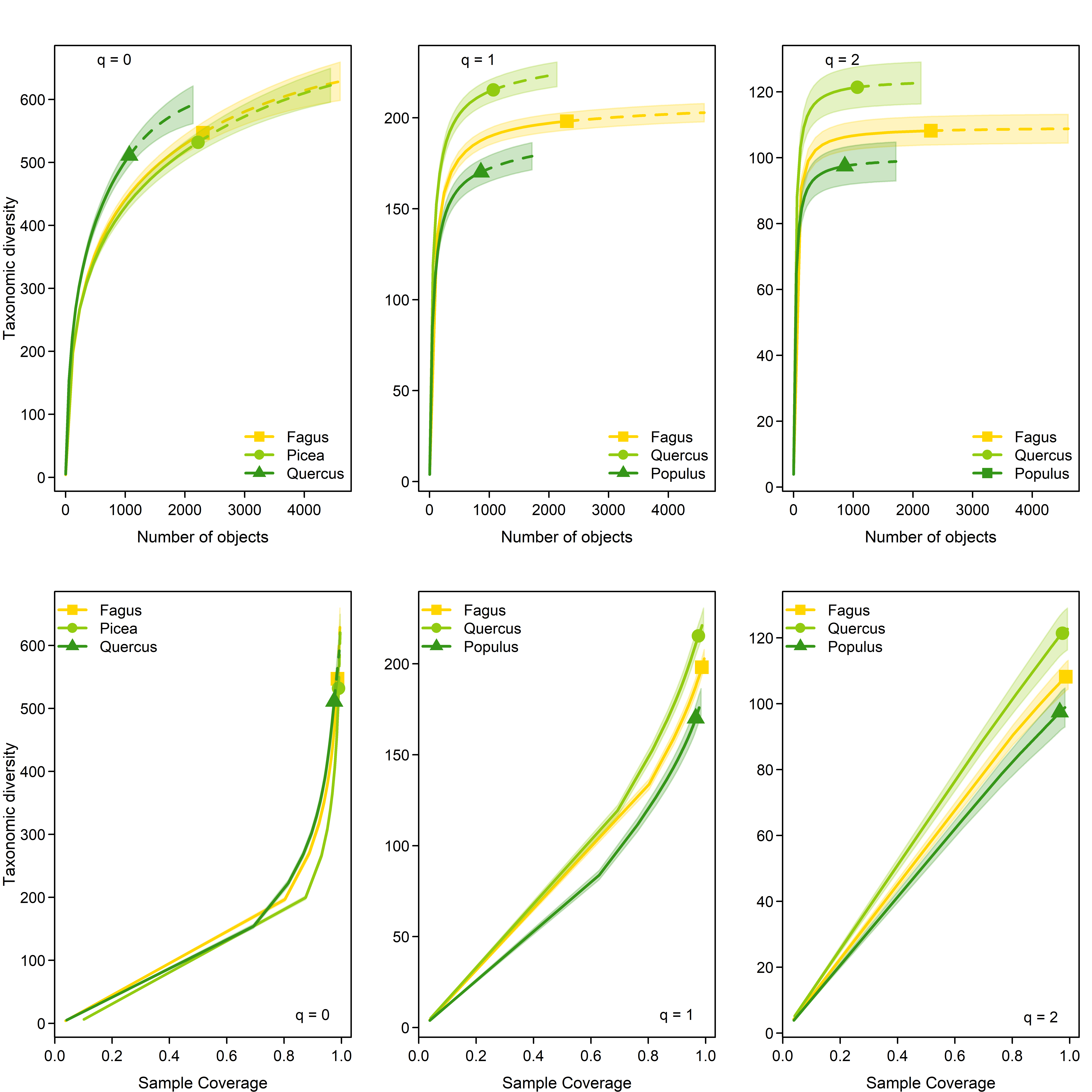

Supplement Figure S7: **Assessment of the sample-size-based and coverage-based rarefaction and extrapolation curves for diversity (Hill numbers) of orders q = 0, 1 and 2.** The panels in the top row display the sample-size-based rarefaction and extrapolation curves up to double the reference sample size. For common (q = 1) and dominant (q =2) beetle species, the sampling curves (top row, middle and left panel) tend to stabilize, implying that the corresponding coverage-based sampling curves can be extrapolated to complete coverage (100 %) to attain the asymptotic estimates. The panels in the bottom row display coverage-based rarefaction and extrapolation curves for diversity up to complete coverage for q = 1 (middle panel) and 2 (right panel), but to a limited coverage value for q = 0 (left panel).

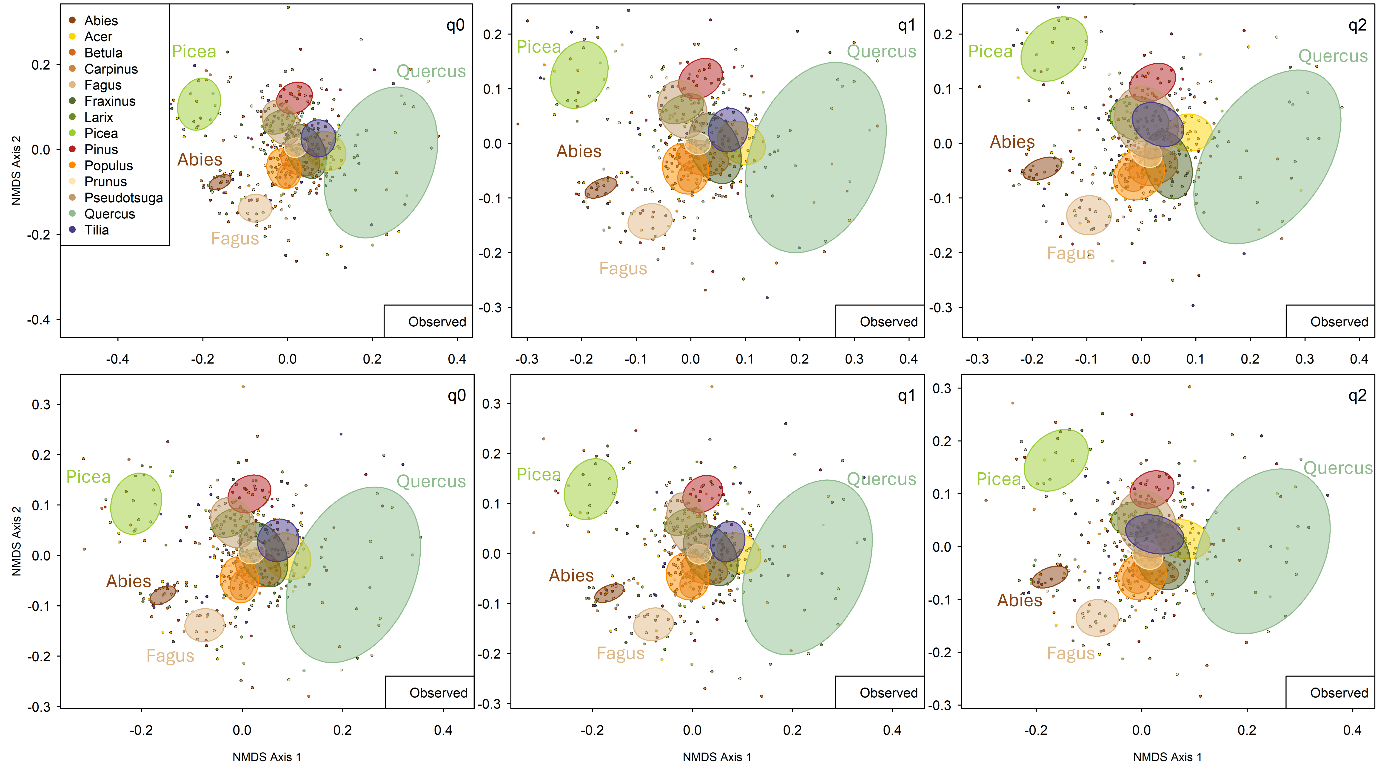

Supplement Figure S8: **Differences in community composition of saproxylic beetles along the Hill numbers**. Nonmetric Multidimensional Scaling (NMDS) for observed data. NMDS indicates the ordinal space within the first two axes of 435 objects, per tree genus. Results are shown for two different subsets of data, where 435 objects per tree genus were randomly selected.

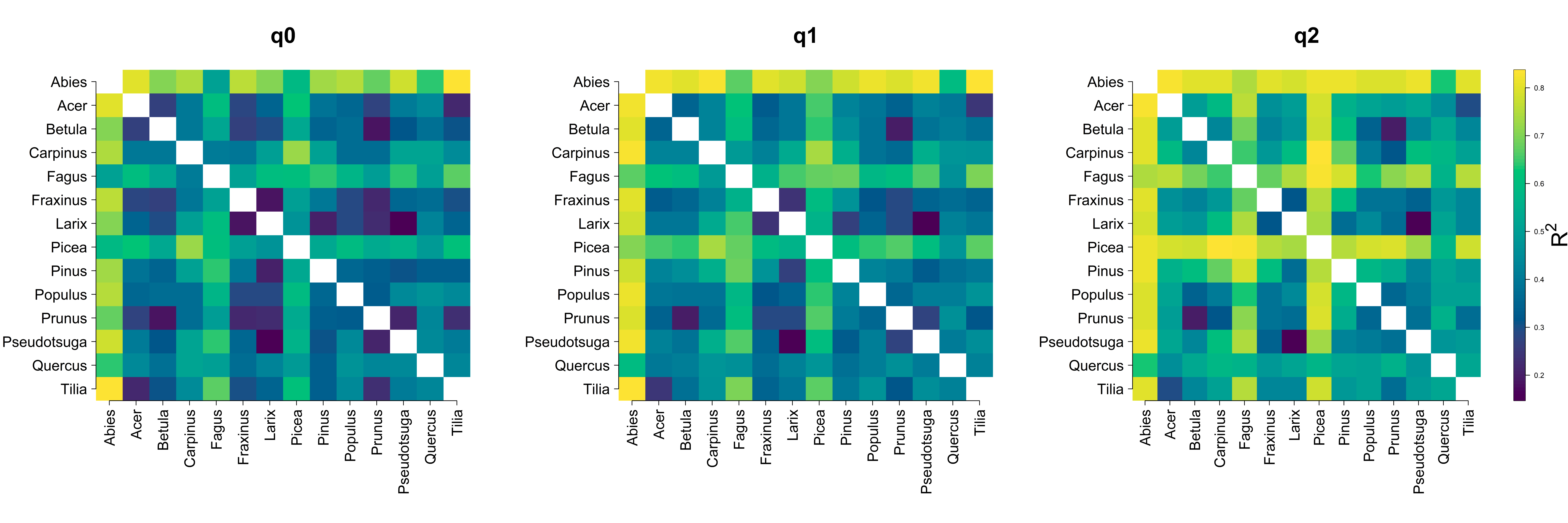

Supplement Figure S9: **Heatmap of pairwise PERMANOVA R² values** showing differences in community composition among 14 tree genera along the Hill numbers. Brighter colours and larger R^2^-values indicate larger compositional differences between genera.

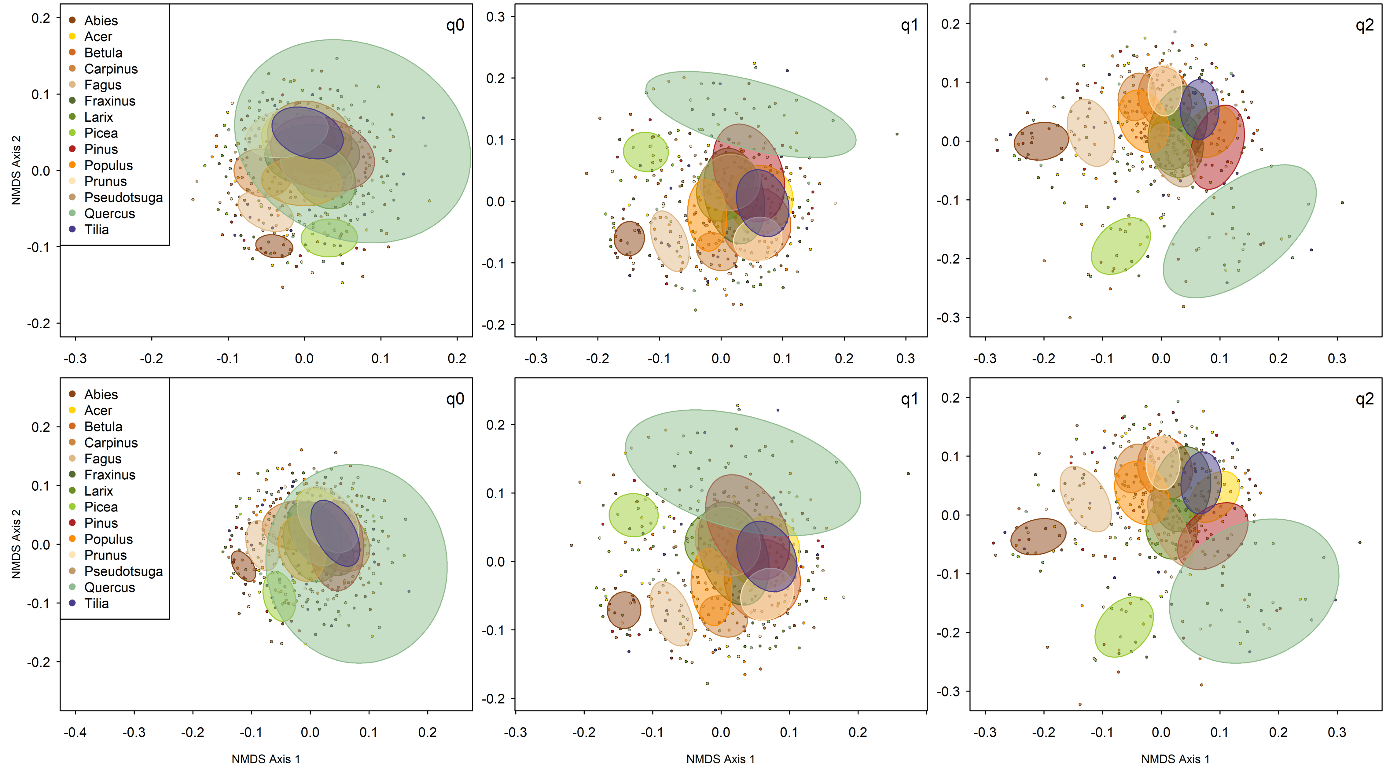

Supplement Figure S10: **Differences in community compositions of saproxylic beetles along the Hill numbers.** Nonmetric Multidimensional Scaling (NMDS) was performed for a standardized coverage of C = 0.97. NMDS indicates the ordinal space within the first two axes of 435 objects, per tree genus. Shown for 2 different subsets of the data.

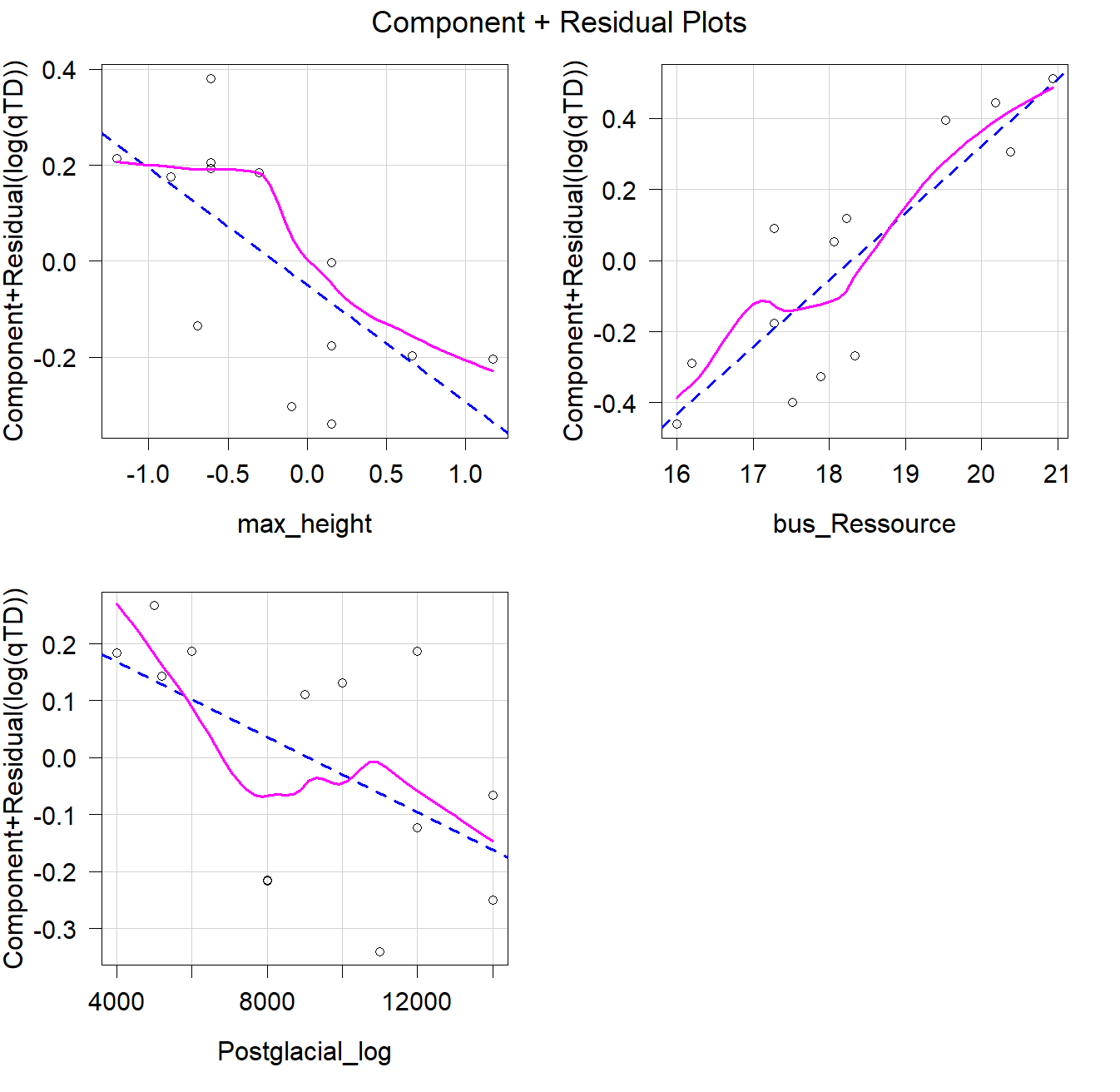

Supplement Figure 11: **Component-plus-residual plots.** Showing the relationship between each continuous predictor (maximum tree height, resource availability and postglacial occurrence) and the response variable (log(qTD), for observed species richness (q = 0)) after adjusting for the effects of the other predictors in the final linear model. The fitted smooth line was inspected to evaluate the assumption of linearity.
